# Trypanosome infection rate and blood-feeding patterns of tsetse fly in an area with a recent human African trypanosomiasis outbreak in Southwest Ethiopia

**DOI:** 10.1101/2024.07.10.602836

**Authors:** Tarekegn Desta, Kokeb Kore, Nigau Eligo, Girum Tamiru, Yilikal Tesfaye, Bernt Lindtjørn, Fekadu Massebo

## Abstract

**Introduction:** The recent outbreak of Human African Trypanosomiasis (HAT) in the Deme River Valley, Ethiopia, has highlighted the need for more comprehensive data on the infection rate of trypanosomes and the feeding patterns of tsetse. The study aimed to assess the infection rate of trypanosomes and the feeding patterns of tsetse flies collected using various attractants.

**Methodology/principal findings:** Traps with various bait odors were placed randomly and rotated monthly. Anesthetized and recently deceased tsetse flies were examined to determine the infection rate using microscopy, and the origins of the blood meal were determined using the polymerase chain reaction technique. We used the Poisson regression model to analyze count data and the binary logistic regression model to assess binary outcomes (positive or negative) and infection rates. Of the 1208 tsetse flies captured, 751 (62.2%) were *Glossina pallidipes*, while 457 (37.8%) were *Glossina fuscipes fuscipes*. The trypanosome infection rate in tsetse flies was 17.1% (95% CI: 14.2-20.3). Of the 103 positive tsetse flies, 66% were infected with *T. congolense*, 24.3% with *T. vivax*, 1.9% with *T. brucei*, and 7.8% with either *T. congolense* or *T. brucei* (the immature stages identified in the mid-gut). The infection rate of trypanosomes in *G. pallidipes* (19. 6%) was higher than that of *G. f. fuscipes* (12.8%). Traps using a combination of cow urine and acetone captured tsetse flies with a higher infection rate of trypanosomes (21.2%), followed by acetone alone (18.9%), cow urine alone (12.2%), and traps without bait (6.1%). Of 107 freshly fed tsetse flies, 23.4% were fed to dogs, 8.4% to humans, 7.5% to cattle, and 4.7% to goats, including the mixed blood meals.

**Conclusions/main suggestions:** The high trypanosome infection rate in tsetse flies indicates an increased risk of trypanosomiasis infection. Although tsetse flies seem attracted to dogs, the potential risk of human exposure must also be considered. Further research is needed to understand the role of dogs in parasite transmission.

**Authors’ summary:** African Animal Trypanosomosis (AAT) is a significant issue in animal production in sub-Saharan African countries, including Ethiopia. The recent outbreak of HAT in the Deme River Valley, Ethiopia, has emphasized the need for more comprehensive data on the infection rate of trypanosomes and the feeding patterns of tsetse flies to understand human exposure. The current study revealed a high infection rate of trypanosomes in tsetse flies, with *T. congolense* having the highest prevalence, followed by *T. vivax*. Public concern is related to the *T. brucei* complex, as it is possibly the strain that causes HAT. Combining cow urine with acetone could significantly improve tsetse trapping and be used in tsetse suppression programs to reduce the risks of infectious tsetse flies. Analysis of the blood meal origins revealed that tsetse flies feed on various vertebrate animals, including dogs and humans. Urgent research is needed to understand the role of dogs in parasite transmission. It is recommended that large-scale tsetse control be implemented using baits such as fermented cow urine in combination with acetone for tsetse suppression.

## Introduction

African Animal Trypanosomosis (AAT) is a major problem in animal production in sub-Saharan African countries, including Ethiopia [1]. The disease also poses a public health threat in areas where African Human Trypanosomiasis (AHT) is common [2]. It is caused by protozoan parasites and transmitted cyclically by the tsetse fly (*Glossina* spp.). The parasites affect vertebrate animals such as livestock, wildlife, and humans [2,3]. According to the World Health Organization, more than 55 million people in Sub-Saharan Africa are at risk of HAT infection [3]. Around 30,000 new sleeping sickness cases were reported. In Ethiopia, around 180,000 to 200,000km² of agricultural land, 14 million cattle and small ruminants, roughly 7 million equines, and 1.8 million camels are at risk of AAT [4]. In 1967, HAT was first identified in Ethiopia’s Baro and Akobo River valleys and later in the southern part of the Omo River basin, and the pathogen was identified as *Trypanosoma brucei rhodesiense* (*T. b. rhodesiense*)[5,6].

In 2022, fifty-five years after the initial epidemic in western Ethiopia, a type of sleeping sickness known as *T.b. rhodesiense* caused an outbreak in the Deme River Valley. This posed a threat to local populations, leading to public health concerns and resulting in loss of lives [7]. This suggests that tsetse flies can bite humans and wild and domestic animals in the region, but it’s still unknown which hosts the flies choose for their blood meals. Additionally, it’s uncertain whether the parasites that cause sleeping sickness in humans are consistently present (enzootic) or occur irregularly (sporadic). Sleeping sickness is a zoonotic disease, meaning it can be transmitted from animals to humans, and human activities such as encroaching into forests for firewood collection, expanding agriculture, and resettling areas can facilitate the spread of the disease [8].

In Ethiopia, there are five species of tsetse flies: *Glossina morsitans submorsitans*, *Glossina pallidipes*, *Glossina fuscipes fuscipes*, *Glossina tachinoides*, and *Glossina longipennis* [4]. The most economically important species is *G. pallidipes*, followed by *G. f. fuscipes*. HAT and AAT are primarily vectored by *G. pallidipes* and *G. f. fuscipes* [9]. There are also five trypanosome species that cause animal trypanosomiasis, namely *Trypanosoma congolense*, *Trypanosoma vivax*, *Trypanosoma brucei brucei*, *Trypanosoma evansi*, and *Trypanosoma equiperdum* [4]. Additionally, *T.b. rhodesiense* is the pathogen that causes HAT [7].

Tsetse flies prefer feeding wild game animals to domestic ones. However, if there is a shortage of wild game, they may start feeding on domestic animals, especially cattle [10]. Identifying the sources of blood meals is crucial for locating tsetse fly hosts, which could be involved in the transmission of parasites and efforts to manage them. The survival of the zoonotic form of trypanosome parasites depends on both domestic and wild animals [11]. To effectively control zoonotic HAT, it is important to identify the sources of the blood meals. Finding trypanosomes in tsetse species indicates exposure to human and animal parasites and is essential for understanding the epidemiology of trypanosomiasis. Furthermore, determining the risk of trypanosomiasis in humans and domestic livestock is crucial for developing effective control measures. Therefore, having more information on the rates of trypanosome infection in *Glossina* species and the specific trypanosome species involved is vital for better understanding the epidemiology and developing efficient control techniques.

Tsetse fly control is now being carried out through various methods, including insecticide spraying, insecticide-impregnated traps and targets, and insecticide-free traps [12]. These traps attract tsetse flies visually, and their efficacy can be increased using different olfactory baits. Understanding the effective odor baits, feeding patterns, and trypanosome infection rates of different tsetse fly species is crucial to controlling trypanosomiasis in Ethiopia effectively. Therefore, this study aimed to estimate the trypanosome infestation and blood-feeding patterns of tsetse flies using different odor bait collections.

## Materials and methods

### Study area description

The study was conducted in southwestern Ethiopia in the Deme River Valley, Gamo Zone. The Deme River basin contains several small and large streams. The largest river in the area is the Deme River, which feeds the Kulano River. At Dana *Kebele*, the Gogara River meets the Deme River, and the combined rivers flow into the Omo River Valley (Figure 1). Another more minor tributary of the Deme River, the Kurtume River, joins the Deme River in Sikole *Kebele*. These water bodies create an optimal environment for the multiplication of tsetse flies.

**Figure 1:**
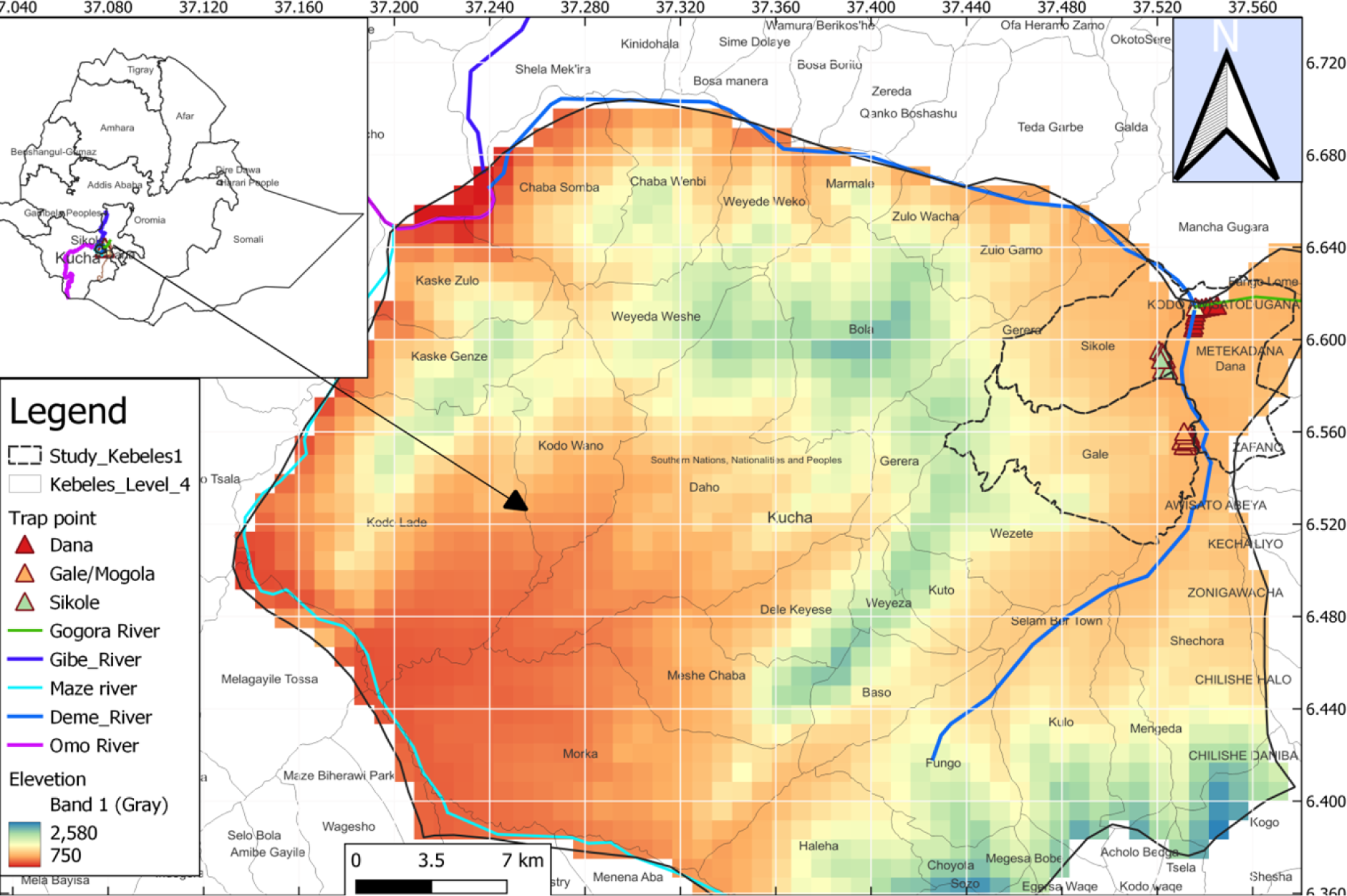
Map of the study area.

The study area has a sub-humid climate with moderately high temperatures and an average yearly rainfall ranging from 900mm to 1800mm [13]. The dry season stretches from December to March, while the wet season is from April to November. The highest recorded temperature was 32.8°C. The cooler months are June, July, and August, with the lowest temperature being 17.5°C. The area is home to wild animals, including warthogs, bush pigs, monkeys, hyenas, gazelles, and rabbits. Domestic animals found here include cattle, goats, sheep, and dogs. It has been reported that there are two types of tsetse flies in the Deme River Valley - *G. pallidipes* and *G. f. fuscipes* [5,14].

AAT is a major challenge in the valley, and several programs have been initiated to control tsetse fly populations. The programs involved removing shrubs that were believed to be tsetse fly habitats. In 1997, the Southern Tsetse Eradication Project (STEP) initiative launched an organized tsetse control plan that introduced various techniques such as insecticide-impregnated targets, ground spray, aerosol, and cattle pour-on using synthetic pyrethroids [14]. Every two to three months, 1% deltamethrin was applied to the backs of cattle. Sterile males (SIT) of two species of tsetse fly (*G. pallidipes* and *G. f. fuscipes*) have been released by aircraft every week for seven years since 2011. During 2011-2014, pour-on and other insecticide-based methods for controlling tsetse fly populations were halted due to concerns that insecticide-treated animals might kill the sterile male tsetse flies released into the field. This interruption led to a significant increase in trypanosome infection in cattle. Farmers in the area have expressed dissatisfaction with the renewed tsetse fly suppression activities following the rise in tsetse flies.

Animals can be exposed to tsetse fly bites when they graze on grasslands, bushes, and water sources where the flies are active. HAT was documented in the same valley in 2022 [7]. People at risk of exposure included daily laborers working in agriculture investment near the Deme River, firewood collectors, herders, and new settlement residents near the valley.

### Study design and period

A longitudinal entomological study was conducted from September 2023 to February 2024 to collect tsetse flies using baited Nguruman (NGU) traps. The sampling occurred over two consecutive days each month, with the attractant locations being rotated monthly. Ten NGU traps were placed in selected locations daily in each Kebele, resulting in twenty monthly traps. Three of the ten traps were baited with acetone. Another three traps were baited with cow urine that had been fermented for three weeks. In addition, three traps were baited with a combination of cow urine and acetone in separate bottles, while one trap was installed without any odor bait. The feet of the traps’ central and lateral poles were greased to deter predators. Fly sampling took place for six months, and 360 traps were deployed in the three selected Kebeles.

We performed monthly randomization and rotation of odor baits to minimize variation due to trap location. This was achieved by creating ten lottery cards with codes representing odor baits. Upon arriving at a trap location, a card was randomly drawn, and the corresponding odor was used. Each position had an equal chance of receiving any odor in the initial round. However, in subsequent rounds, the odor bait used in the previous month was excluded from the draw for the same position to avoid the repetition of the same odor in a given location.

The geographic location of the traps was determined using the Global Positioning System (GPS), which allowed us to calculate the distance between traps and deployment sites. The distance between traps within each site ranged from 200 to 250m. Meanwhile, the distance between the Dana and Gale sites was 6.8km, while between the Sikole and Dana sites was 3.2km, and between the Sikole and Gale sites was 4.1km.

### Tsetse fly sampling

Ten *Kebeles* located near the Deme River Valley were infested with tsetse flies. Three *Kebeles* (Gale, Sikole, and Dana) were randomly selected for the study. To capture the tsetse flies, NGU traps were placed in the tree sheds, which were their resting locations. The trap deployment was carried out at 8:00 am and collected at 6:00 pm. The trapping sites were kept open to allow the flies to see the traps. When there was no open location to deploy the traps, the vegetation obstructing trap visibility was cut within a 3–6-meter radius around the trap, depending on the height of the vegetation. The traps were hung 20 cm above the ground. We set up traps in different locations, such as riverbanks, forests, farmland, and areas where people live. Each trap caught flies that were labeled in the field. We collected the remaining flies in a cage, which we then placed in an icebox. We used these flies for species identification and dissection and to identify where they got their blood meals from.

### Tsetse identification

The tsetse control personnel training manual key for *Glossina* species identification was used to distinguish between different species of tsetse flies [15]. Adult Tsetse flies possess unique morphological traits that differentiate them from other insects. These features include a distinct proboscis, folded wings at rest with hatchet cells, and branched arista hairs on their antennae. *Glossina* species can be identified based on crucial characteristics such as the color of their hind and front legs’ tarsal segments and the color and form of the dorsal surfaces of their abdomen, with or without banding. It’s possible to distinguish male flies from females by examining their abdomen. The male fly has a hypopygium on the posterior end of the ventral abdomen, which is absent from the underside of a female tsetse [15].

### Tsetse dissection and parasite identification

All live and recently deceased non-teneral flies were selected without bias during the dissection of tsetse flies. The trap cages were kept in an icebox during transportation from the field to the dissecting room, and each cage was handled individually. To verify that they were non-teneral, the flies were removed from the cage, and gentle pressure was applied to their thorax to euthanize them. The location, odor bait type, and geographic coordinates from the cage’s label were documented on Excel datasheets. The species and sex of each fly were determined. Non-teneral flies were identified based on their abdomen and the absence of a dark or brown color, which indicated recent blood intake.

To ensure accurate results, dissections were conducted on freshly euthanized or recently deceased flies. This was important because trypanosomes are less likely to be found in dead, desiccated flies. Initially, a drop of saline solution was placed in the center of a petri dish, and the fly was positioned on its back on top of it to prevent dehydration and maintain the isotonic state of the dissected organs. Using fine watchmaker forceps, each wing and leg was carefully removed from the fly, ensuring other organs were not damaged. The organs where trypanosomes develop, including the mid-gut, salivary glands, and proboscis, were dissected and inspected under a microscope at a magnification 40x.

The process was carried out in a specific sequence to prevent contamination caused by trypanosomes found in the midgut during dissection. First, the proboscis was dissected, followed by the salivary glands, and finally, the midgut. To start, the proboscis was firmly held on the thecal bulb and removed with the help of fine forceps while being observed through a dissecting microscope at low magnification (6x).

The labrum, hypopharynx, and labium were separated with the tip of tiny forceps and a hypodermic needle. The dissected organ was placed in one drop of saline solution on the glass slide and covered with a cover slip. The fly’s thorax was held with one pair of fine forceps, the base of the abdomen was grasped with a second pair of fine forceps, and the abdomen was gently pulled back so that the “skin” slowly tears while the gut and other internal contents stretch out but do not break. Salivary glands on each lateral side of the abdomen were picked from the lateral sides of the first abdominal segments of the fly using a pair of fine forceps and then placed on a glass slide with a drop of 0.9% standard saline solution and covered with a cover slip. The abdominal contents were squeezed or pulled from the skin, and the mid-gut was picked up and placed on a glass slide with a drop of saline solution on one side, covered by a cover slip. To avoid contamination of the sample by dissecting needles and forceps, the needles and forceps were wiped with clean cloths, rinsed with sterile distilled water, and immersed in saline solution between dissections of each organ and fly.

After dissecting all three organs of a fly, the dissected components were viewed using a compound microscope at 40x magnification. The existence of trypanosomes was determined by the motility in the places (organs) of the fly where trypanosome parasites were isolated due to their life cycle. Thus, infection in the labrum or hypopharynx alone (the proboscis) was classified as ‘vivax-type.’ Infections in the proboscis and mid-gut were classified as ‘congolense-type,’ while infections of the proboscis, mid-gut, and salivary glands were classified as ‘brucei complex’ infections [16].

### Blood meal source identification

Freshly fed tsetse flies were carefully collected and stored in individual Eppendorf tubes. The tubes were placed in a plastic zip bag with silica gel beads and transported to the Advanced Medical Entomology and Vector Control laboratory at Arba Minch University. Upon arrival, the specimens were kept in a deep freeze at - 80°C. When it was time to extract DNA for blood meal detection, the flies were crushed, and the Cetyltrimethylammonium bromide (CTAB) DNA isolation procedure was used[17].

The samples were analyzed to identify blood meal sources by multiplex PCR method using primers from humans, dogs, cattle, goats, sheep, pigs, and horses. Host species were inferred based on the DNA band size.

### Statistical analysis

This study examined two main outcomes: the infection status of tsetse flies with trypanosome parasites (a binary outcome), and the number of tsetse flies collected per trap per day. To analyze the infection status, we used a binary logistic regression model, while we employed a Poisson regression model to study the number of flies collected per trap per day. We estimated the parameters for both models using maximum likelihood estimation (MLE) and calculated standard errors to gauge the statistical significance of the factors involved. Model fit was assessed using the Akaike Information Criterion (AIC) and other relevant measures. All the analyses were carried out using R statistical software (version 4.1.1), and we used the ggplot2 package for creating plots.

## Results

### Tsetse fly species in the study sites using various odor baits

The traps in the three *Kebeles* (120 traps in each) captured 1208 tsetse flies, averaging 3.4 per trap per day. The two identified species of tsetse flies were G*. pallidipes* and *G. f. fuscipes*. Out of the 1208 tsetse flies captured, 751 (62.2%) were *G. pallidipes*, with an average catch of 2.1 flies per trap per day. The remaining 457 (37.8%) were *G. f. fuscipes*, with an average catch of 1.3 flies per trap per day. Table 1 summarizes the number of tsetse flies captured in the three study *Kebeles* using various odor baits.

**Table 1:**
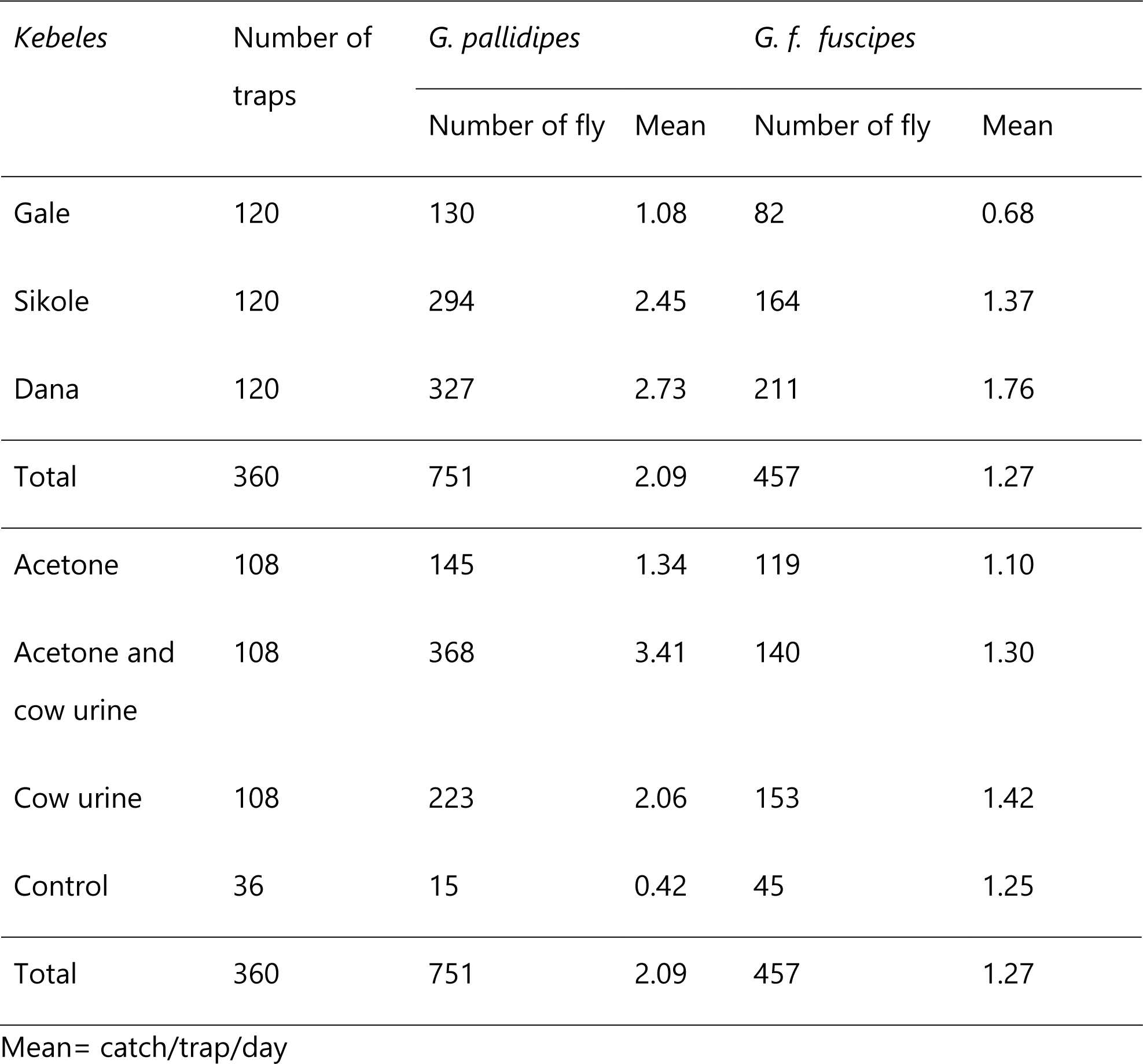
Number of *G. pallidipes* and *G. f. fuscipes* in the study *Kebeles* using various odor baits in Deme River Valley, south Ethiopia.

### Efficiency of odor baits in capturing tsetse flies

In Dana *Kebele*, the number of tsetse flies captured was higher using all odor-baited traps and the control trap. Conversely, all odor-baited traps recorded the lowest number of tsetse flies in Gale Kebele (Figure 2). The number of tsetse flies captured varied significantly among different odor baits and study *Kebeles*. All odor baits captured significantly more tsetse flies than the control, and both Dana and Sikole *Kebeles* had more tsetse fly counts than Gale (Table 2).

**Figure 2:**
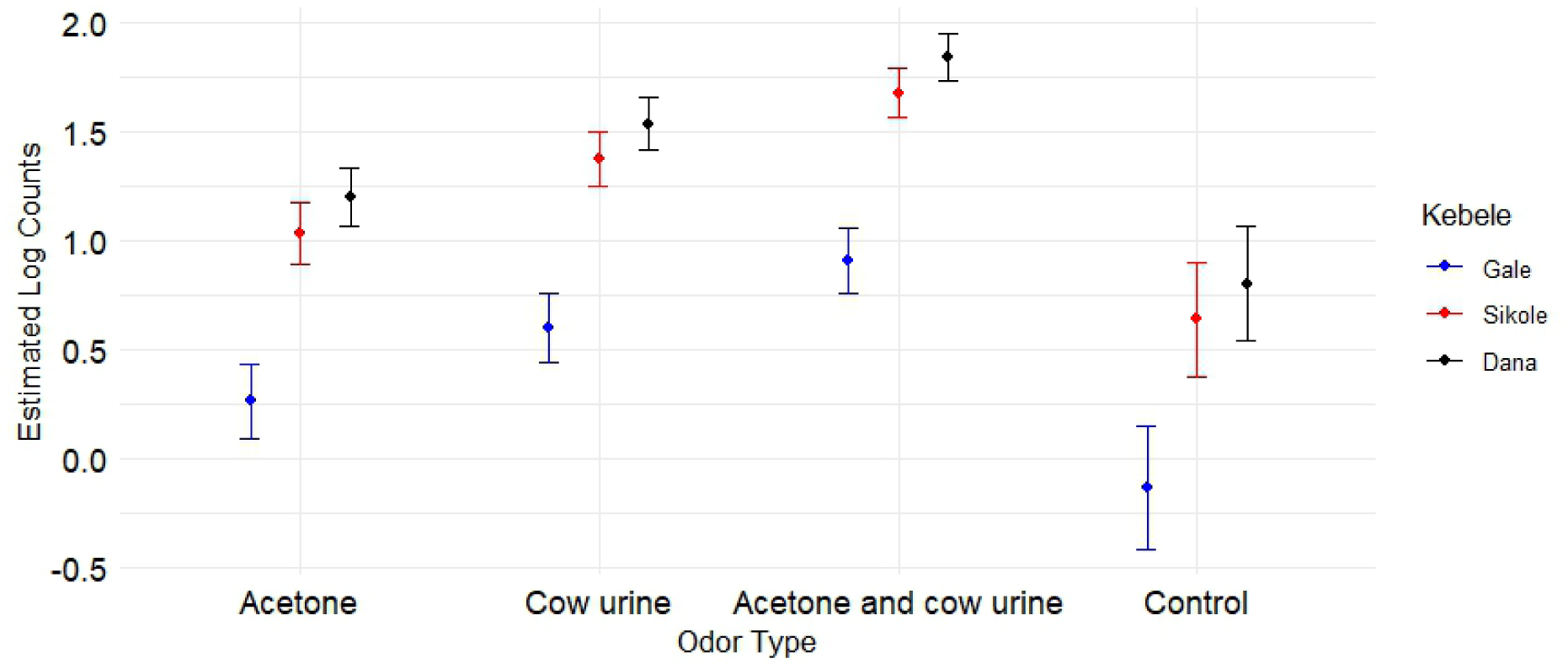
The estimated log counts of tsetse flies in the three study *Kebeles* using various odor baits in the Deme River Valley in Gamo zone, southwest Ethiopia.

**Table 2.** The estimated Log count variation of tsetse flies between odors baits and study *Kebeles* in the Deme River Valley in Gamo zone, southwest Ethiopia.

| Predictors | Categories | Estimate | Std. error | Statistic | P value |
| --- | --- | --- | --- | --- | --- |
| Intercept |  | 0.2636 | 0.0874 | 3.017 | 0.00255 |
| Odor types | Cow urine | 0.337 | 0.0801 | 4.206 | <0.0002 |
|  | Acetone and cow urine | 0.6413 | 0.0756 | 8.481 | <0.0001 |
|  | Control | -0.394 | 0.1429 | -2.76 | 0.0347 |
|  | Acetone (ref) |  |  |  |  |
| *Kebeles* | Sikole | 0.7681 | 0.0831 | 9.243 | <0.0001 |
|  | Dana | 0.9331 | 0.0811 | 11.51 | <0.0001 |
|  | Gale (ref) |  |  |  |  |

Using odor baits improved the effectiveness of NGU traps in catching tsetse flies. A combination of cow urine and acetone was the most effective for catching tsetse flies, followed by cow urine alone. The traps without odor baits caught fewer tsetse flies. The trapping effectiveness of cow urine was significantly higher than that of acetone but significantly lower than that of traps baited with a combination of acetone and cow urine. All traps baited with attractants were more efficient than the control in catching tsetse flies (Figure 3).

**Figure 3:**
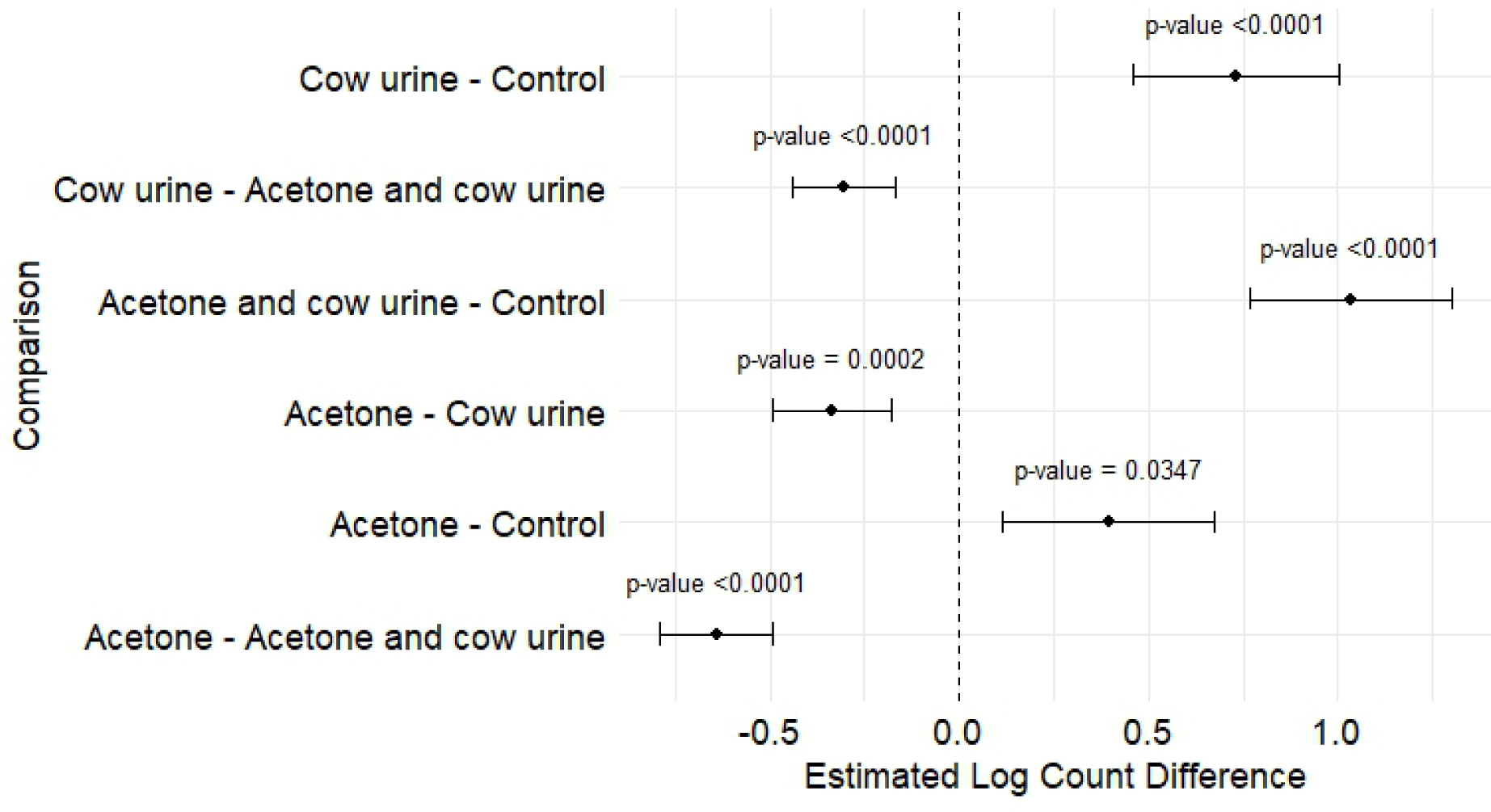
Pairwise comparison of the efficacy of odor baits using log counts of tstse flies in Deme River Valley in Gamo Zone, Southwest Ethiopia.

### The trypanosome infection rate of tsetse flies

Out of a sample of 603 live and recently died non-teneral tsetse flies dissected to determine the trypanosome infection rate, 103 tsetse flies were infected with at least one species of trypanosomes, giving an overall infection rate of 17.1% (one sample proportion 95% CI: 14.2-20.3) and of the 377 dissected *G. pallidipes*, 73 (19.4%; 95% CI 15.6 -23.8) had trypanosome infection in their proboscis, salivary glands, mid-gut, or all of these organs. Likewise, out of the 226 dissected *G. f. fuscipes*, 30 (13.27%; 95% CI: 9.27-18.56) were found to have trypanosome infection in their trypanosome-developing organs. Throughout the study, *G. pallidipes* tested positive for trypanosomes in all *Kebeles* more frequently than *G. f. fuscipes*. Although the infection rate of trypanosomes in *G. pallidipes* was higher than that of *G. f. fuscipes*, the difference was not statistically significant (Figure 4).

**Figure 4:**
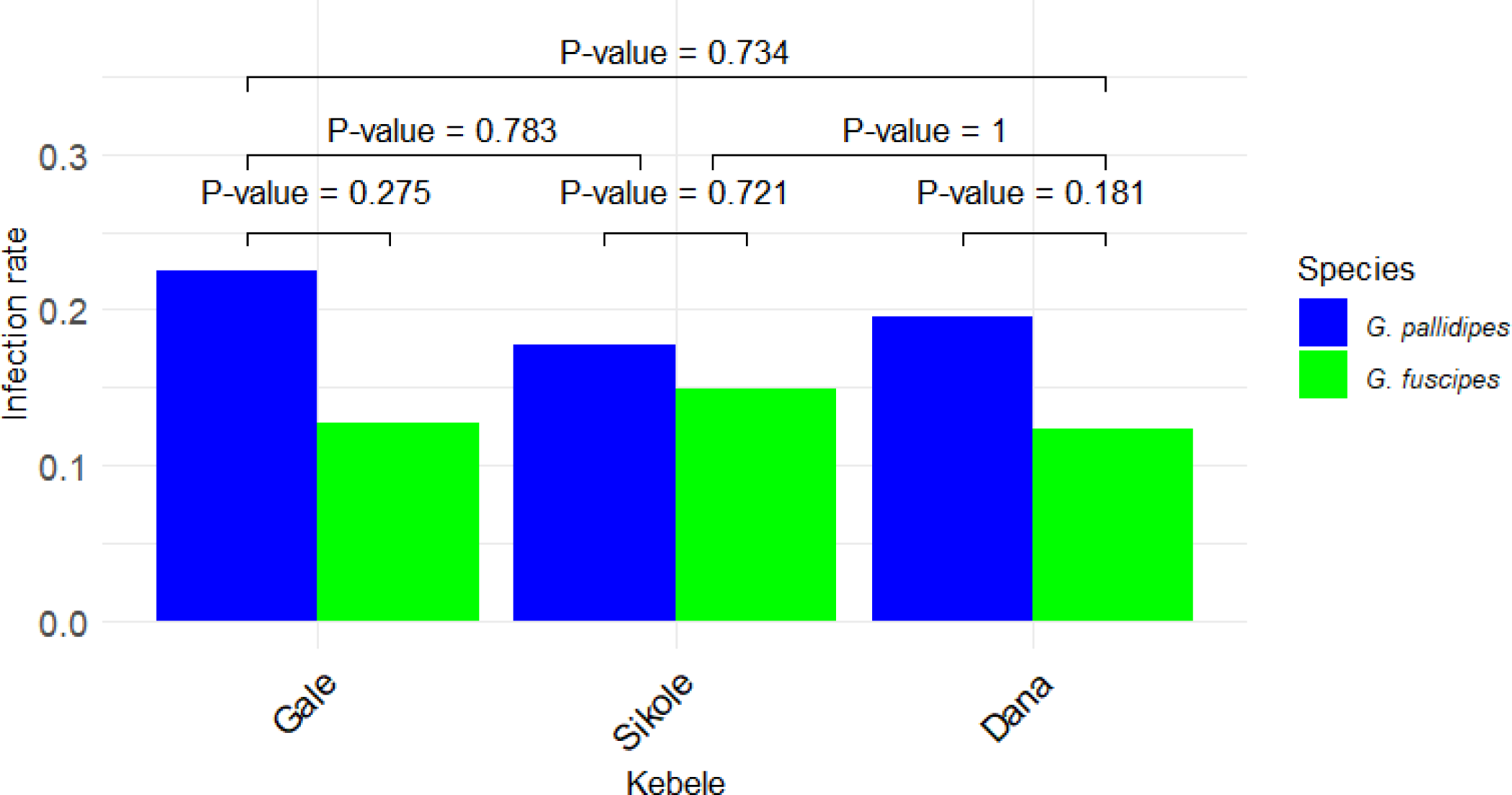
The number of trypanosome-positive tsetse flies over the study period and *Kebeles* using various odor baits in Deme River Valley in Gamo Zone, Southwest Ethiopia.

### Trypanosome species composition in tsetse flies

Out of the 103 positive tsetse flies, 68 flies (66%) were infected with *T. congolense*, 25 flies (24.3%) were infected with *T. vivax*, and two flies (1.9%) were infected with *T. brucei.* Eight flies (7.8%) were infected with either *T. congolense* or *T. brucei* (the immature stages identified in the mid-gut). These trypanosomes were isolated from their respective development sites in the tsetse flies. *Trypanosome congolense* and *T. vivax* were the two most common parasites found during the study period; however, *T. brucei* was a rarely documented species in the tsetse flies (Figure 5).

**Figure 5:**
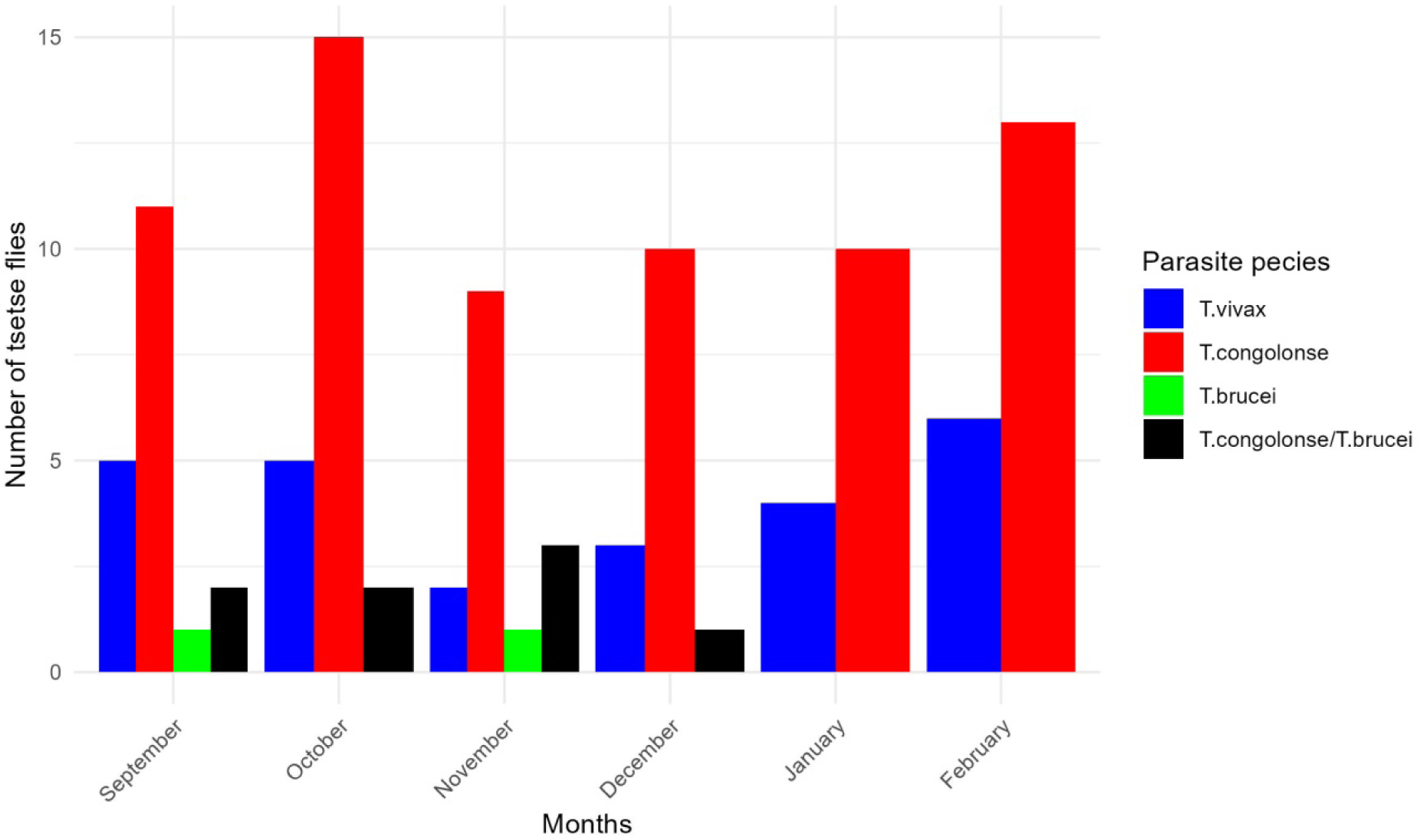
The number of tsetse flies positive for parasite species over the study period in the Deme River Valley in Gamo Zone, Southwest Ethiopia.

Out of the 377 *G. pallidipes* dissected, 18 tested positive for *T. vivax*, resulting in an infection rate of 4.8%. Seven of the 226 *G. f. fuscipes* dissected tested positive for *T. vivax*, resulting in an infection rate of 3.1%. The infection rate of *T. congolense* in *G. pallidipes* was 13.3% (50 out of the 377 dissected), while in *G. f. fuscipes*, the infection rate was 7.9% (18 out of the 226 dissected). Two *G. f. fuscipes* tested positive for *T. brucei* complex. Among the eight tsetse flies that tested positive for either *T. congolense* /*T. brucei* complex (immature identified in the mid-gut of tsetse flies), five were identified as *G. pallidipes*, and three were identified as *G. f. fuscipes*. Of the three *Trypanosoma* species identified in the study area, *T. congolense,* and *T. vivax* were found in all *Kebeles* during the study period (Figure 6).

**Figure 6:**
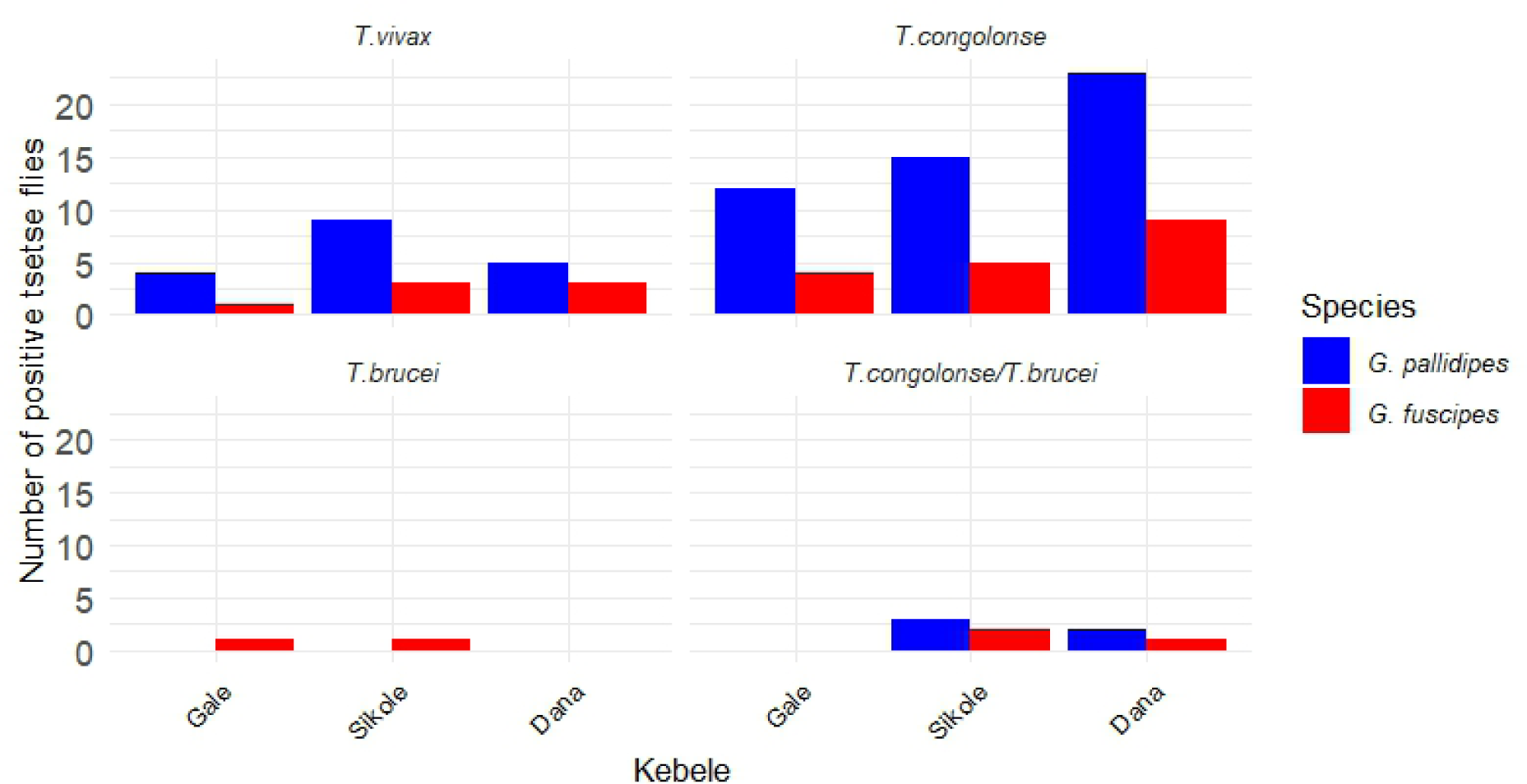
The number of positive tsetse flies by month, odor type, *Trypanosoma* species, and in the study *Kebeles* in the Deme River Valley in Gamo Zone, Southwest Ethiopia.

The infection rate of trypanosome was highest in tsetse flies captured by traps baited with a combination of cow urine and acetone (21.4%; 55/257), followed by those baited with only acetone (19.8%; 25/126) and cow urine alone (11.2%; 21/187). Tsetse flies captured in traps without odor baits (6.1%; 2/33) had the lowest trypanosome infection rate.

Although the prevalence of infection rates of tsetse flies captured by various odors seems to vary, there was no statistically significant difference between odors, tsetse fly species, and Kebeles (Table 3). For example, there was no significant difference in infection likelihood between tsetse flies attracted by a combination of acetone and cow urine compared to flies attracted by acetone alone (*P* – value = 0.883).

**Table 3.** The estimated infection rate of tsetse flies between odors baits, study *Kebeles* and tsetse species in the Deme River Valley in Gamo zone, southwest Ethiopia.

| Predictors | Categories | Estimate | Std. error | Statistic | P. value |
| --- | --- | --- | --- | --- | --- |
| Intercept |  | -1.195 | 0.327 | -3.66 | <0.001 |
| Odor types | Cow urine | -0.682 | 0.323 | -2.11 | 0.034 |
|  | Acetone and cow urine | 0.04 | 0.273 | 0.146 | 0.883 |
|  | Control | -1.238 | 0.767 | -1.61 | 0.106 |
|  | Acetone(ref) |  |  |  |  |
| Species | *G. fuscipes* | -0.343 | 0.243 | -1.42 | 0.156 |
|  | *G. pallidipes* (ref) |  |  |  |  |
| Kebele | Sikole | -0.071 | 0.30 | -0.23 | 0.814 |
|  | Dana | -0.079 | 0.293 | -0.27 | 0.785 |
|  | Gale(ref) |  |  |  |  |

### Blood meal sources of tsetse flies

Blood meal source identification was conducted on 107 freshly fed tsetse flies. The results showed that 31.8% (34/107) of the tsetse flies tested positive for one or more tested vertebrate animals DNA, while 68.2% (73/107) tested negative for the tested vertebrate animals. None of the freshly fed tsetse flies were positive for sheep, horses, and pig blood meal origins.

Overall, tsetse flies most commonly feed on dogs (*Canis lupus*) (14), followed by cattle (6), humans (*Homo sapiens*) (2), and least frequently on goats (*Caprines*) (1) (Table 4).

**Table 4:**
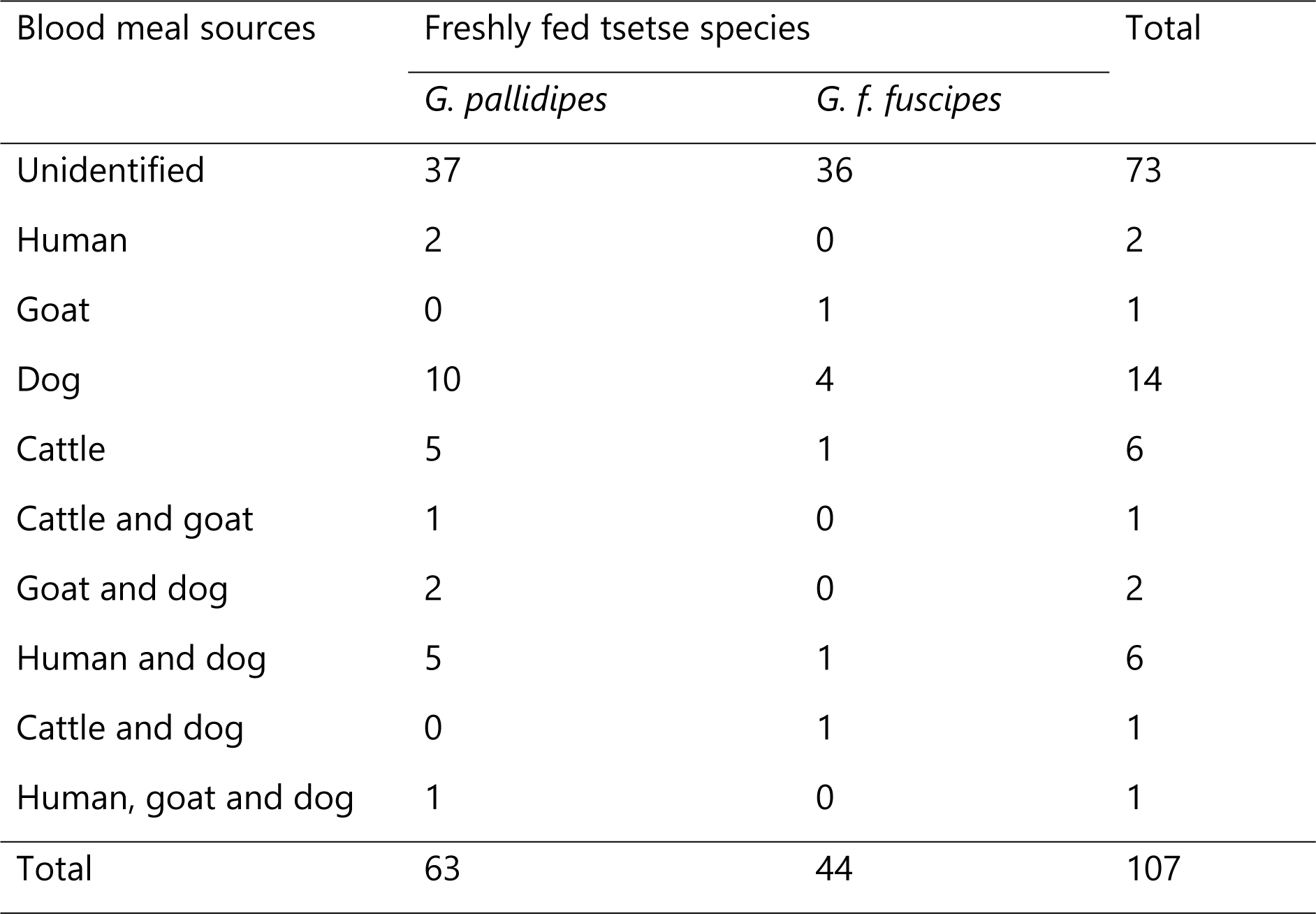
Blood meal sources of tsetse fly blood meal sources in Deme River Valley, Gamo zone, south Ethiopia.

| Blood meal sources | Freshly fed tsetse species |  | Total |
| --- | --- | --- | --- |
|  | *G. pallidipes* | *G. f. fuscipes* |  |
| Unidentified | 37 | 36 | 73 |
| Human | 2 | 0 | 2 |
| Goat | 0 | 1 | 1 |
| Dog | 10 | 4 | 14 |
| Cattle | 5 | 1 | 6 |
| Cattle and goat | 1 | 0 | 1 |
| Goat and dog | 2 | 0 | 2 |
| Human and dog | 5 | 1 | 6 |
| Cattle and dog | 0 | 1 | 1 |
| Human, goat and dog | 1 | 0 | 1 |
| Total | 63 | 44 | 107 |

## Discussion

The trypanosome infection rate was higher in tsetse flies, especially in *G. pallidipes,* compared to *G. f. fuscipes*. The most effective baits for catching tsetse flies were a combination of cow urine and acetone, followed by traps with cow urine alone. Traps without baits had the lowest catch rate of tsetse flies. The traps using a combination of cow urine and acetone captured tsetse flies with a higher infection rate of trypanosomes, followed by acetone alone, cow urine alone, and traps without bait. *T. congolense* and *T. vivax* were identified as the most common parasites in the study area. Tsetse flies feed on various vertebrate animals, such as dogs, cows, and humans, and often have mixed blood meals from different animals.

Two species of tsetse flies, *G. pallidipes* and *G. fuscipes*, were found in the study area. These two species are among the five *Glossina* species found in Ethiopia, particularly in the South Regional State [4]. Both species were recently documented in the Dame River Valley [18]. There is a high risk of tsetse fly invasion from the Omo River Valley into the Dame River Valley, and without adequate control, re-invasion is also likely. To prevent the invasion, insecticide-impregnated barrier targets were placed at the entrances of the tsetse fly from the Omo River Valley to the Deme River Valley. However, tsetse control activities and maintenance of barrier targets were halted for the past six years, allowing different species of tsetse flies to re-invade the area [14]. The current high density of the two species of tsetse flies in the study valley could have resulted from the interrupted control tools. The proportion of *G. pallidipes* was higher than that of *G. fuscipes*, possibly due to the use of the NGU trap. *G. pallidipes* prefer a horizontal orientation facilitated by the NGU trap, while *G. fuscipes* prefer a vertical orientation. Another reason for the difference in species proportion may be *G. f. fuscipes’* poor response to the odor attractants used.

The study compared the infection rates of trypanosomes in two species of tsetse flies. The results showed that *G. pallidipes* had a higher trypanosome infection rate than *G. f. fuscipes*. This outcome aligns with previous research conducted in Gojeb Valley, Southwest Ethiopia [9], and the Baro Akobo River system, western Ethiopia [19]. However, the trypanosome infection rate in the current study site was lower than in previous Nech Sar National Park research on the same species, *G. pallidipes* [20]. This variation could be attributed to the differences in the detection tools used for trypanosome infection in tsetse flies. Rodrigues et al. [20] used the highly sensitive PCR assay, while we used microscopy.

The finding also indicated that *T. congolonse* was the most prevalent, followed by *T. vivax*, while *T. brucei* complex was the least prevalent in both tsetse fly species. The highest prevalence of *T. congolense* could be due to its high maturation rate of the parasites and the vector’s high susceptibility to this parasite species [21]. Even though the prevalence of *T. brucei* complex is low, it suggests the risks of HAT as the parasite is likely to be *T. b. rhodesiense*, the causative agent of HAT.

The higher infection rate in tsetse flies caught in traps containing a mixture of cow urine and acetone may be due to the attraction of older tsetse flies to this odor combination. The age of tsetse flies is the most critical factor in determining the likelihood of trypanosome infection, as older flies are more likely to harbor mature trypanosome parasites [16]. Trypanosome infection prevents tsetse flies from successfully taking a blood meal and can cause interruptions during their meal intake [22]. Additionally, the parasites compete for proline, a vital energy source for tsetse flies [23]. Therefore, infected tsetse flies, needing energy, are more likely to roam around odor-baited traps than uninfected ones, making them more susceptible to being caught or killed by these devices. Consequently, using odor-baited traps can reduce the risk of infectious tsetse flies and help in their control.

Odor-baited NGU traps outperformed traps without odor baits for catching *G. pallidipes*. Previous studies have also shown that traps with odor baits perform better [24]. The traps baited with cow urine and acetone, followed by cow urine alone, caught the most *G. pallidipes*, while traps without bait caught the least. When these baits are used with killing agents, the optimized odor bait combinations can be used as a “push-pull” deployment strategy for area-wide tsetse fly control [25]. Large-scale deployment of odor-baited traps could be possible for effective tsetse control in a shorter period. In practical terms, using cow urine to catch tsetse flies is crucial as it emphasizes its practicality and cost-effectiveness as an odor attractant in tsetse control strategies. Cow urine can be easily obtained and effective, mainly after it has been left to stand for up to three weeks. It is well-suited for community-based African trypanosomiasis control using traps, especially in resource-poor countries. Our research indicates that using cow urine alone or combined with baited traps could result in significant fly catches for controlling *G. pallidipes*.

The PCR-based blood meal origins detection results indicate that tsetse flies feed on various animals, such as dogs, cattle, humans, and goats. Despite the small sample size, dogs were identified as the most common blood meal source, followed by cattle and humans. The blood meal origins of tsetse flies vary, possibly due to the availability of host species and the ecology and species of tsetse fly vectors [25, 26]. Human activities such as hunting with dogs, farming and gathering firewood in the forest, and visiting rivers and streams can expose humans and dogs to tsetse fly bites. Unlike in wildlife-dominated regions where tsetse flies tend to feed on a single preferred host species, the multiple vertebrate hosts’ feeding patterns of tsetse flies were observed [26]. The diverse blood-feeding patterns of tsetse flies could be due to the absence of preferred hosts, potentially leading to higher exposure to various parasites.

### Conclusions

The high infection rate of trypanosomes in tsetse flies in the area indicates a high risk of AAT. The presence of *T. congolense* in tsetse flies suggests an increased risk of AAT. Although the prevalence of *T. brucei* complex in tsetse flies was low, it could pose a public health threat as two strains cause HAT. Combining cow urine and acetone could significantly improve tsetse trapping and be used in tsetse suppression programs to reduce the risks of infectious tsetse flies. Analysis of tsetse fly blood meals indicates that they feed on various vertebrate animals, including dogs and humans. This finding suggests that dogs may play a role in spreading African Trypanosomosis as they were the primary source of blood meals for tsetse flies. Further research is urgently needed to understand the role of dogs in parasite transmission and to implement large-scale tsetse control using baits such as fermented cow urine and its combination with acetone for tsetse suppression.

## Acknowledgments

We acknowledge the field team involved in setting and transporting traps.

## Author contributions

Conceived the study: TD, FM

Sample collection: TD, KK

Involved in the laboratory work: TD, KK

Conducted molecular laboratory work: TD, NE, GT

Statistical data analysis: YT, TD, FM

Supervise the laboratory work: FM, KK

Drafted the manuscript: FM

Critically commented on the manuscript and approved: TD, KK, NE, GT, YT, BL, FM

## Ethical approval

Not applicable

